# Analysis of SARS-CoV-2 genomes from across Africa reveals potentially clinically relevant mutations

**DOI:** 10.1101/2020.09.08.287201

**Authors:** Modeline N. Longjohn, Olivia S. Egbule, Samuel O. Danso, Eugene E. Akujuru, Victor T. Ibubeleye, Christabel I. Oweredaba, Theodora Ogharanduku, Alexander Manu, Benson C. Iweriebor

## Abstract

SARS-CoV-2 is a betacoronavirus, the etiologic agent of the novel Coronavirus disease 2019 (COVID-19). The World Health Organization officially declared COVID-19 as a pandemic in March 2020 after the outbreak in Wuhan, China, in late 2019. Across the continents and specifically in Africa, all index cases were travel-related. Understanding how the virus’s transportation across continents and different climatic conditions affect the genetic composition and the consequent effects on transmissibility, infectivity, and virulence of the virus is critical. Thus, it is crucial to compare COVID-19 genome sequences from the African continent with sequences from selected COVID-19 hotspots/countries in Asia, Europe, North and South America and Oceania.

To identify possible distinguishing mutations in the African SARS-CoV-2 genomes compared to those from these selected countries, we conducted *in silico* analyses and comparisons. Complete African SARS-CoV-2 genomes deposited in GISAID and NCBI databases as of June 2020 were downloaded and aligned with genomes from Wuhan, China and other SARS-CoV-2 hotspots. Using phylogenetic analysis and amino acid sequence alignments of the spike and replicase (NSP12) proteins, we searched for possible vaccine coverage targets or potential therapeutic agents. Identity plots for the alignments were created with BioEdit software and the phylogenetic analyses with the MEGA X software.

Our results showed mutations in the spike and replicate proteins of the SARS-Cov-2 virus. Phylogenetic tree analyses demonstrated variability across the various regions/countries in Africa as there were different clades in the viral proteins. However, a substantial proportion of these mutations (90%) were similar to those described in all the other settings, including the Wuhan strain. There were, however, novel mutations in the genomes of the circulating strains of the virus in African. To the best of our knowledge, this is the first study reporting these findings from Africa. However, these findings’ implications on symptomatic or asymptomatic manifestations, progression to severe disease and case fatality for those affected, and the cross efficacy of vaccines developed from other settings when applied in Africa are unknown.

## Introduction

Severe Acute Respiratory Syndrome Coronavirus 2 (SARS-CoV-2) is the etiologic agent for the so-called Coronavirus disease 2019 (COVID-19). COVID-19 first emerged in Hubei Province, China, in December 2019; the second coronavirus to have arisen from China in the last two decades^1,2^. Within months, SARS-CoV-2 spread rapidly both nationally and internationally, resulting in global outbreaks of pneumonia-like symptom clusters and death in some cases. Consequently, the World Health Organization publicly declared COVID-19 a pandemic on 11th March 2020. Besides SARS-CoV-2, four strains of coronavirus pathogenic to humans are associated with clinical symptoms, including common cold and pneumonia^3^. Two other coronaviruses, SARS-CoV (responsible for the Severe acute respiratory syndrome – SARS) and MERS (which causes the Middle Eastern respiratory syndrome) were notable for causing infections with higher fatalities deaths^4,5^. However, outbreaks caused by SARS-CoV and MERS (though still circulating) were relatively limited compared to those caused by SARS-CoV-2^6^.

The Severe Acute Respiratory Syndrome Coronavirus 2 (SARS-CoV-2) is a novel beta coronavirus (family *Coronaviridae*, genus *Betacoronavirus*, species *Severe acute respiratory syndrome-related coronavirus*)^1,2^. SARS-CoV-2 is an enveloped virus that has a positive sense of single-stranded viral RNA genome of approximately 30kb^7^. The SARS-CoV-2 genome consists of 14 open reading frames (ORFs) preceded by transcriptional regulatory sequences (TRSs), which form transcriptional units8. The largest transcriptional unit includes the ORF1a and b regions, which translates into two precursor polyproteins (PP1a and PP1b), that are subsequently cleaved by viral proteases into functional non-structural proteins (Nsps)^8^. Other ORFs encodes four main structural proteins, including surface spike (S) glycoprotein, envelope (E) protein, membrane (M) protein and nucleocapsid (N) protein (See Fig. 1), each of which has specific functions that dictate SARS-CoV-2 interactions and biology^9–11^.

**Figure 1:**
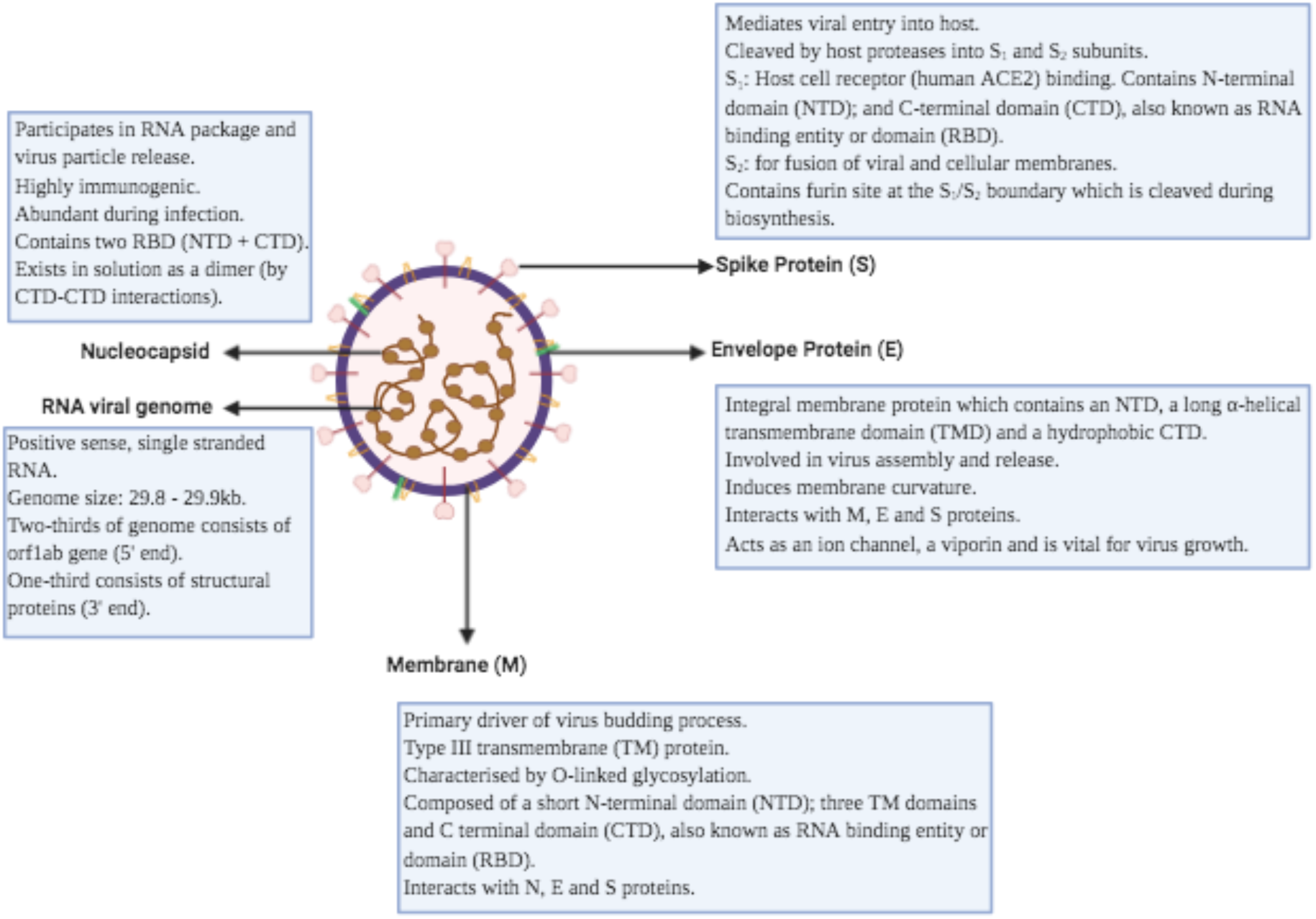
SARS-CoV-2 viral structure showing structural proteins and the RNA viral genome.

**Figure 2:**
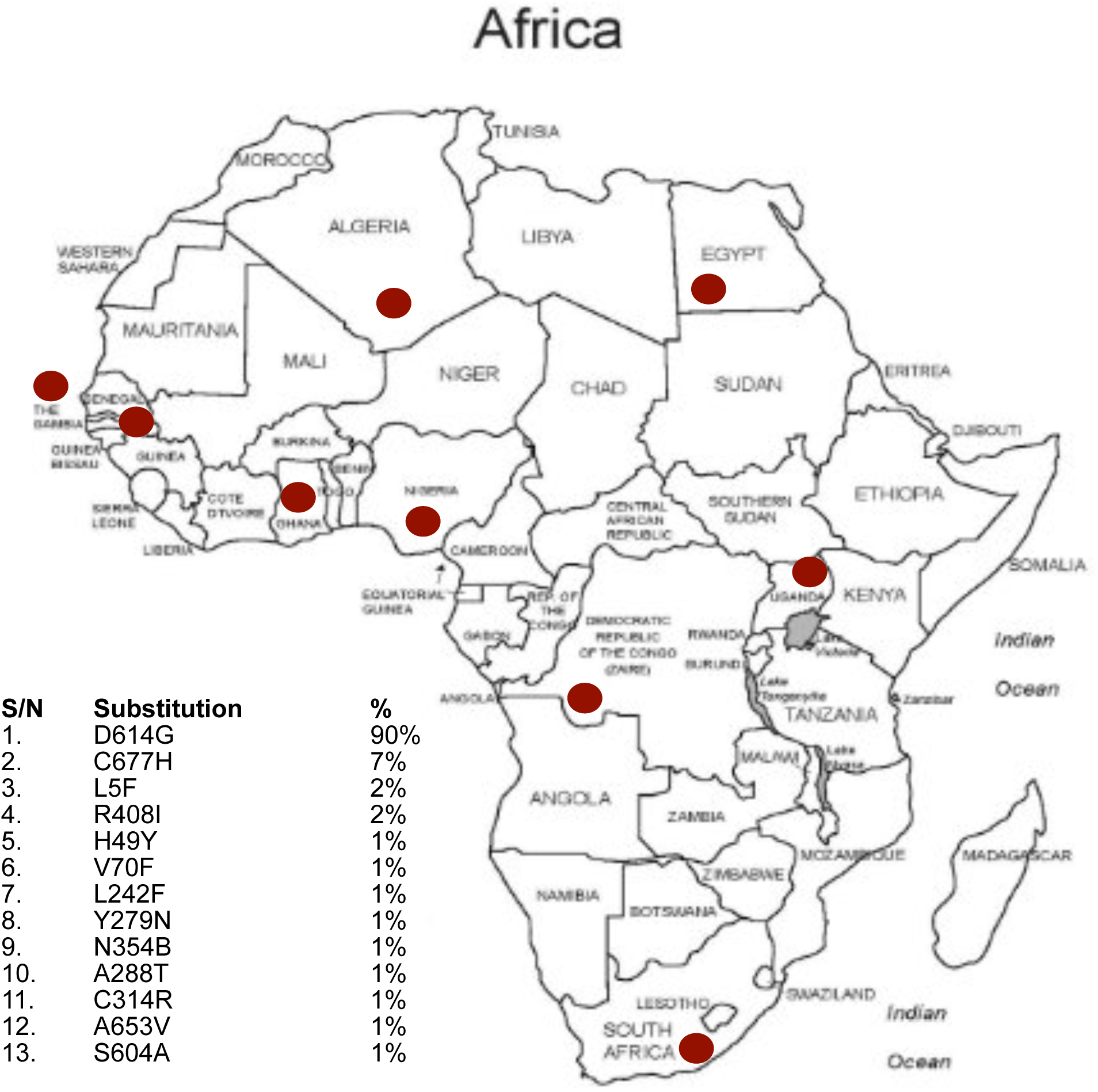
Amino acid substitutions observed in the spike proteins of Coronavirus sequences circulating in Africa. Image created using Affinity designer version 1.8.4.

The pathophysiology and virulence of SARS-CoV-2 are heavily reliant on the function of Nsps and other structural proteins. Specifically, Nsps can prevent host immune response^1^. For structural proteins, the E protein is crucial in viral pathogenicity and promotes viral assembly and release^2^. The S protein also facilitates viral entry into target cells, having an affinity for the angiotensin-converting enzyme 2 (ACE2) receptor expressed on the surface of cells of the lungs, heart, renal system^2,12^. Since the pandemic onset, scientists have contributed over 34,300 genome sequences to the Global Initiative on Sharing Avian Influenza Data (GISAID) and NCBI GenBank databases as of May 2020. Of this, Africa has only contributed 224 genome sequences. This is mostly due to Africa’s reduced scientific research capacity, despite an abundance of expertise. As of 12th May 2020, Africa accounts for about 2% of the global COVID-19 caseload^3^. This has a massive implication for the African continent, especially as the continent currently lacks the health care system capacity to manage a full-blown outbreak effectively.

Currently, there is no known cure for COVID-19, with most pandemic response systems across the globe not being adequate in curbing the spread of the virus. Hence, there is a need to develop a vaccine(s) and drug(s), in which research teams across the globe are in a race against time to develop. However, this process is a multi-step process that relies heavily on specific molecular and pharmacological data. For insights into the molecular basis of developing vaccines and drugs, the pool of genome sequences of SARS-CoV-2 from all over the world is currently being mined and analyzed by researchers. Conversely, many vaccines under development are currently at phase III of trials, implying that the vaccine development may have commenced before the index case was reported in some African countries. This, in turn, raises an important question; would the treatments and vaccines, when available, be useful in Africa?

A previous study showed that residues near some amino acid (Lysine 31, Tyrosine 41, 82-84, and 353-357) in human ACE2 were important for SARS-CoV-2 binding^4^. To investigate mutations in the amino acid residues of SARS-CoV-2 circulating in Africa, we analyzed whole-genome sequences of SARS-CoV-2 from Africa deposited in the GISAID and NCBI GenBank databases. Because all cases in Africa arose from travel, it became compelling to compare COVID-19 sequences generated from the African continent with strains from COVID-19 hotspots Wuhan, China, and other countries across the world’s different regions. We conducted complete genome analyses of viruses from Africa and compared them with reference sequences from Wuhan and other parts of the world using phylogenetic analysis. We also investigated the spike protein’s amino acid sequences and the replicase (NSP12) to identify similarities and differences. This study shows, for the first time, an in-depth analysis of the SARS-CoV-2 molecular epidemiology in Africa and will potentially provide insights into the mutations/variations in relevant proteins of viruses circulating in Africa.

## Materials and Methods

### Search Strategy for Identification of COVID-19 SEQUENCES

We searched the SARS-CoV-2 NCBI (https://www.ncbi.nlm.nih.gov/sars-cov-2/) and GISAID (https://www.gisaid.org/) databases for SARS-CoV-2 genomic sequences from Africa, specific countries within Asia, Oceania, North America and Europe. For our analyses, nucleotide and amino acid sequences of the SARS-CoV-2 genome were used. Also included were spike, NSP12 and ORF1ab nucleotide and protein sequences, respectively.

### Inclusion Criteria

Only complete genomes with at least 29kb of length were selected for the phylogenetic analysis, while ORF1ab and surface glycoproteins with 1273 amino acids were downloaded from the Coronavirus genomes NCBI website. Table 1 shows the details of the curated proteins of novel coronavirus data downloaded. Efforts were made to download proteins for spike and NSP12 from different geographical regions globally, including Africa, Asia, Europe, North America, South America, and Oceania. All curated surface glycoproteins from Africa with omitted or ambiguous amino acid residues were eliminated from the analysis.

**Table 1.** African Countries and the country of Origin of the index case: All initial cases of SARS-CoV-2 in African countries were travel related. The source of the index cases for the sequences used in this study are outlined in the table above.

| Africa Region |  |  |  |
| --- | --- | --- | --- |
| Sub-region | Country (City of index case) | Date of index case | Origin of index case |
| East Africa | Uganda (Entebbe) | 18<sup>th</sup> March 2020 | United Arab Emirates |
| West Africa | Gambia (Banjul) | 17<sup>th</sup> March 2020 | United Kingdom |
|  | Ghana (Accra) | 12<sup>th</sup> March 2020 | Norway via Turkey |
|  | Nigeria (Lagos) | 27<sup>th</sup> February 2020 | Italy |
|  | Senegal (Dakar) | 2<sup>nd</sup> March 2020 | France |
| Central Africa | Democratic Rep of Congo (Kinshasa) | 10<sup>th</sup> March 2020 | France |
| North Africa | Algeria (Bilda) | 17<sup>th</sup> February 2020 | Italy |
|  | Egypt (Cairo) | 14<sup>th</sup> February 2020 | China |
| Southern Africa | South Africa (Johannesburg) | 1<sup>st</sup> March 2020 | Italy via UAE |
| Comparison countries from other regions |  |  |  |
|  | China (Wuhan) |  |  |
|  | Brazil (Rio de Janeiro) |  |  |
|  | Italy (Milan) |  |  |
|  | Spain (Madrid) |  |  |
|  | United Kingdom (London) |  |  |
|  | United States of America (New York) |  |  |
|  | Australia (Sydney) |  |  |
|  | New Zealand |  |  |

### Amino Acid alignment of the surface glycoprotein (spike protein)

All African surface glycoproteins of the SARS-CoV-2 obtained from the NCBI GenBank were aligned using ClustalW and compared to a reference strain from Wuhan, China. After that, a consensus sequence was created for the African sequences and a global consensus sequence for all spike proteins from different geographical regions of the world, respectively. A total of 44 African surface glycoproteins sequences along with a reference strain that were analyzed are as follow: YP_009724390_Wuhan, QJY78020.1,QJY78032.1, QJY78044.1, QJY78056.1, QJY78068.1, QJY78092.1, QJY78104.1, QJY78116.1, QJY78128.1, QJY78140.1, QJY78152.1, QJY78164.1, QJY78176.1, QJZ28114.1, QJZ28126.1, QJZ28347.1, QJZ28359.1, QKE11078.1, QJX45321.1, QJX45333.1, QJX45344.1, QJX45356.1, QJX45380.1QKK12863.1, QKI36913.1, QKD20860.1,QKG82981.1,QKG82993.1,QKF95995.1, QIZ15537.1, EPI_ISL_418241_Algeria, EPI_ISL_418217_Senegal,EPI_ISL_418207_Senegal, EPI_ISL_418217_Senegal, EPI_ISL_421573_SouthAfrica, EPI_ISL_418241_Algeria, EPI_ISL_421574_SouthAfrica, EPI_ISL_428857_Gambia, EPI_ISL_428855_Gambia, EPI_ISL_430819_Egypt, EPI_ISL_437339_DRC, EPI_ISL_447234_DRC, EPI_ISL_413550_Nigeria, EPI_ISL_430820_Egypt.

### Amino acid alignment of the NSP12-Replicase protein

ORF1ab amino acid sequences of SARS-CoV-2 from Africa were downloaded, and 1050 NSP12 sequences were identified. The consensus sequence of the African NSP12 was created and aligned with the Wuhan reference strain, and identity plot was created with BioEdit. Analysis of the NSP12 also performed with global sequences of the replicase protein, and efforts were made to include representatives of the protein from the world’s six geographical regions. The global consensus was created from these sequences as was done with the African sequences, and the consensuses were aligned with the Wuhan strain and identity was plotted with the reference strain as was performed earlier with the African consensus sequence. The 28 replicase proteins from Africa and the reference strain that were analyzed are as follow: YP_009724389.1_Wuhan, QJY78018.1, QJY78030.1,QJY78042.1, QJY78054.1, QJY78066.1, QJY78078.1, QJY78090.1, QJY78102.1, QJY78114.1, QJY78126.1, QJY78162.1, QJY78174.1, QJZ28112.1, QJZ28124.1, QJX45319.1, QJX45331.1, QJX45342.1, QJX45354.1, QJX45366.1, QJX45378.1, QKK12861.1,QKI36911.1,QKD20858.1, QKG82979.1, QKG82991.1,QKF95993.1, QKF96005.1, QIZ15535.1.

### Genomes phylogenetic analysis

For phylogenetic analyses, complete SARS-CoV-2 genomes (a total of 56 sequences from Africa and other regions of the world) were obtained from GenBank. Genomes with at least 29kb were used in the analysis, with bat coronavirus genome used as an outgroup. Phylogenetic tree construction by the neighbor joining method was performed using MEGA X software, with bootstrap values of 1000^5^. The percentage of replicate trees in which the associated taxa clustered together in the bootstrap test (1000 replicates) was indicated next to the branches^6^. The tree was drawn to scale, with branch lengths in the same units as those of the evolutionary distances used to infer the phylogenetic tree. Poisson correction method was used to compute the evolutionary distances considering the units of the number of amino acid substitutions per site^8^. All ambiguous positions were removed for each sequence pair (pairwise deletion option). Evolutionary analyses were conducted in MEGA X software while multiple alignment was performed using MUSCLE alignment as implemented in MEGA^9,10^.

## Results

### Amino acid alignment of the surface glycoprotein

The amino acid sequence alignment of African surface glycoproteins against the Wuhan reference strain showed that sequences were almost identical, except for few positions where non-synonymous substitutions occurred. Thirteen mutations were identified to differ between African and non-African SARS-CoV-2 spike proteins (See Table 2). 90% of the African spike proteins had D614G substitution, while 7% had a C677H mutation. L5F and R408I mutations occurred in 2% of all cases. The remaining mutations nine mutations occurred at a rate of 1%.

Of the identified mutations, the D614G substitution (shown in the amino acid alignment in Figure 2) is the most famous mutation among global SARS-CoV-2 sequences. Analysis of the spike proteins from the different geographical regions of the globe revealed that Asia’s sequences have a predominance of D614G substitution. Also, European sequences have mostly the D614G substitution as well as the Oceania sequences.

There is currently no record of the C677H, V70F, L242F, N354B, A288T, C314R, A653V and S604A mutations in the literature. Therefore, this study shows the existence of these mutations at a low percentage for the first time. However, the L5F, R408I and H49Ymutations have previously been reported in the literature.

**Figure 2.**
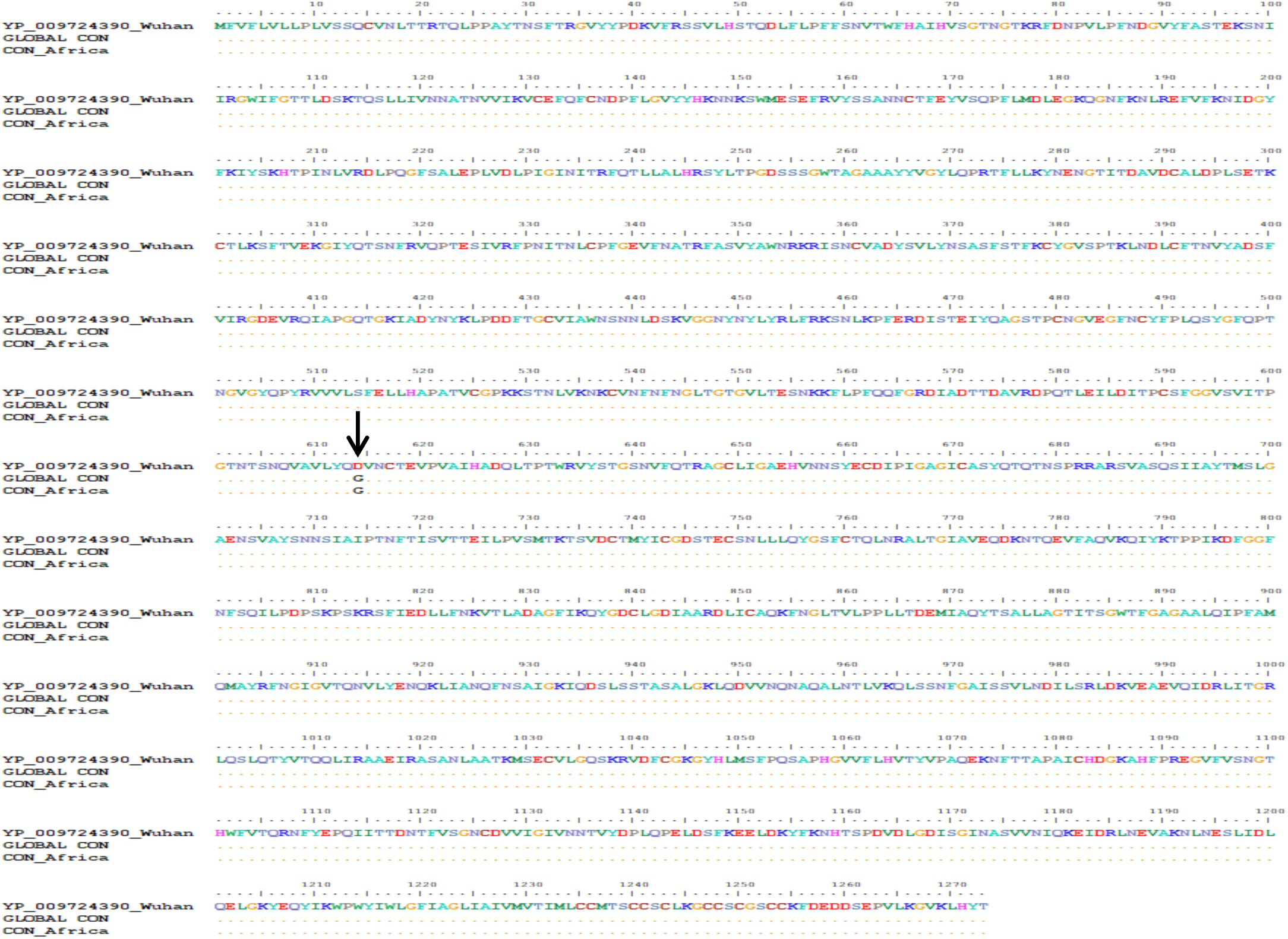
Amino acids alignment of the Spike protein of the novel Coronavirus: Consensus sequences were created from Wuhan, Global and African sequences of spike protein sequences obtained from NCBI GenBank. The consensuses of African and global spike proteins were compared with the reference strain from Wuhan. Dots represent similarity in amino acid, while there was no deletion in the protein. There was an amino acid substitution in the African and global sequences at position D614G, as shown by an arrow.

### Amino Acid alignment of the NSP12-Replicase protein

Analysis of the NSP12 of African, global sequences compared with the Wuhan consensus sequence showed that sequences were almost identical except for deletion at position P314-in both the African and global sequences, as shown by an arrow in Figure 3.

**Fig.3.**
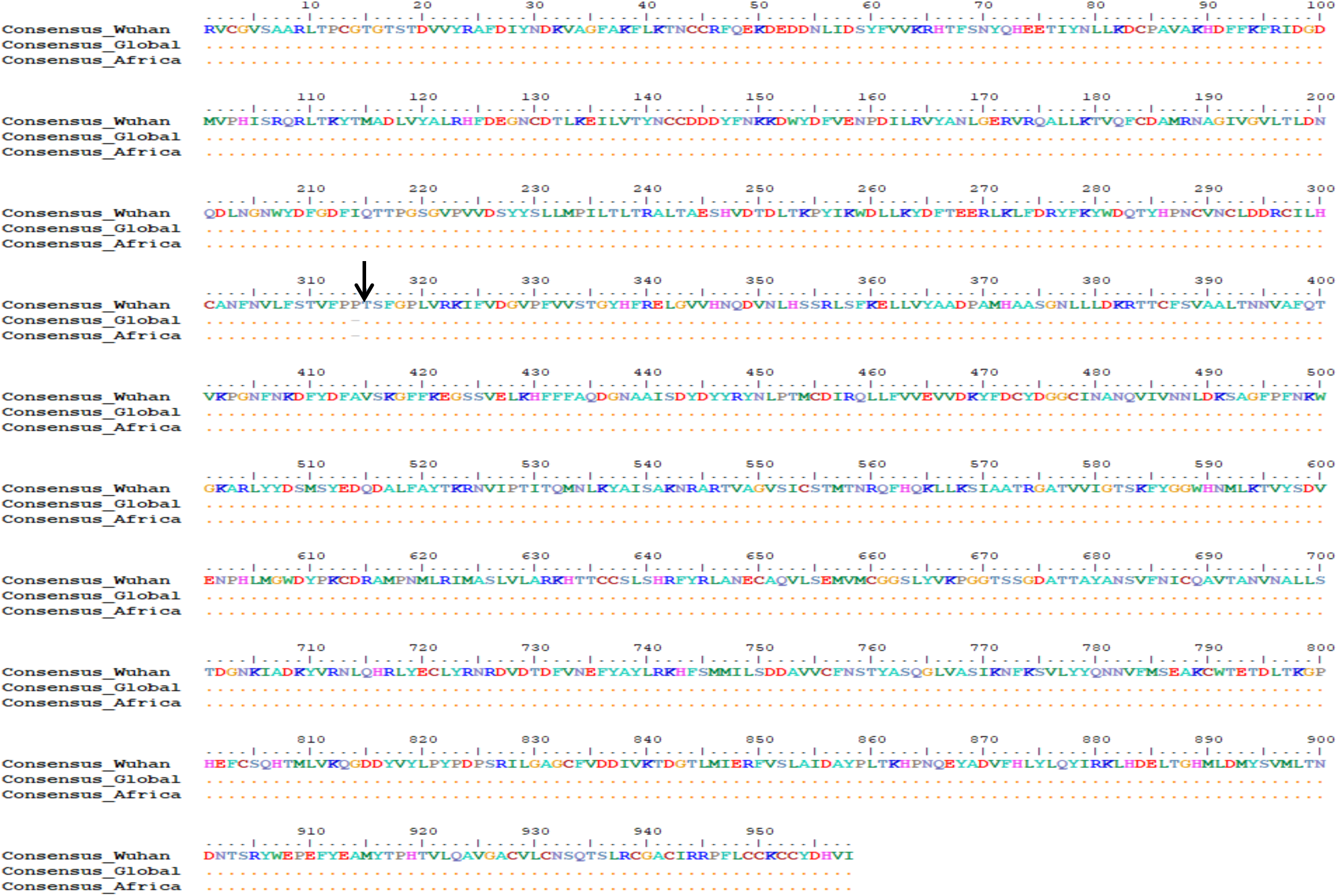
Amino acids alignment of the NSP12 of the novel Coronavirus: Consensus sequences were created from Wuhan, Global and African sequences of NSP12 within the OR1ab sequences obtained from NCBI GenBank. The consensuses of African and global were compared with the consensus generated from Wuhan ORF1ab sequences. Dots represent similarity in amino acid while the dash denotes a deletion. There was a deletion in the African and global sequences at position P314- as indicated by an arrow.

### Phylogenetic analysis of genomes of SARS-CoV-2 from Africa

Phylogenetic analysis of trees based on amino acid sequences of SARS-CoV-2 from different parts of Africa and global reference strains with bat coronavirus sequence showed that sequences originating from African countries clustered with each other overall African sequences did not form a monophyletic cluster as shown in Fig 4. However, there seemed to be multiple strains across South Africa, which did not all align together. This may likely mean that there were multiple introductions of SARS-CoV-2 strains from different regions. Furthermore, the DRC and Algeria also have two different strains, while all other African countries have only one strain. Overall, there seem to be multiple African clades that demonstrate local transmission within the African countries.

**Fig 4.**
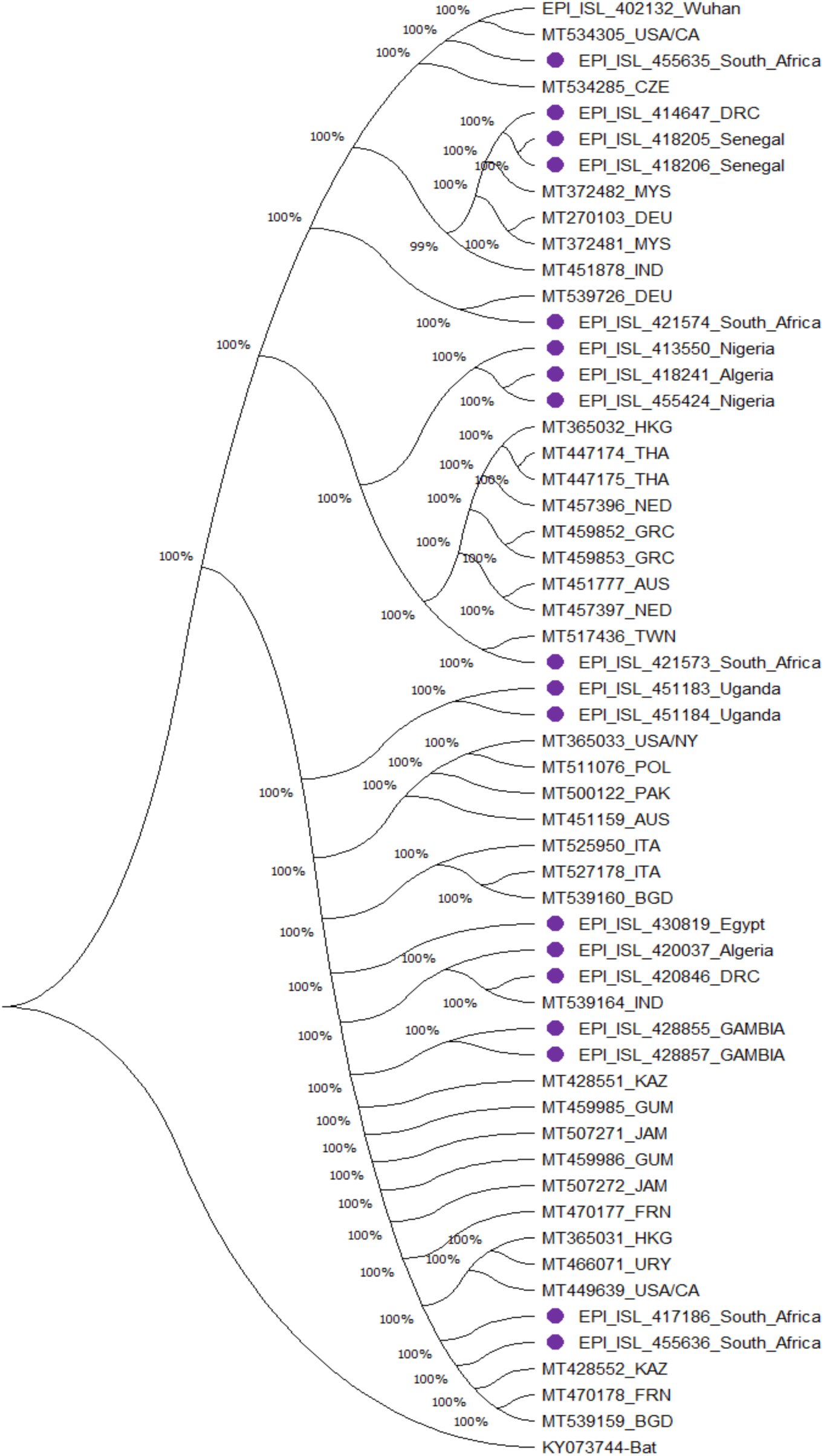
Evolutionary relationships of SARS-CoV-2 genome taxa. The evolutionary relationship between African SARS-CoV-2 sequence in comparison to reference sequences from Wuhan, China, Asia, Australia, Europe and the Americas were inferred using the Neighbor-Joining method. The analysis involved 56 nucleotide sequences. Codon positions included were 1st+2nd+3rd+Noncoding. All ambiguous positions were removed for each sequence pair. There was a total of 27873 positions in the final dataset. Evolutionary analyses were conducted in MEGA X [4]. African sequences are denoted in purple colored balls and abbreviation of the name of country while sequences from Non-African countries were denoted using abbreviation of the country name. This figure shows that the SARS-CoV-2 strain in Africa are evolutionarily related to strain in different countries.

The number of African SARS-CoV-2 complete genomes found at the time of collecting this data (May 31st, 2020) was two hundred and thirteen (213). These sequences were selected to be included in the tree based on the completeness of the genomes.

## Discussion

This is one of the first studies from Africa that describes the genomic comparison of SARS-CoV-2 viruses circulating in Africa, how they compare with the rest of the world and genomic evolutionary trends on the continent. Our results show mutations in the spike protein and replicase proteins of the SARS-CoV-2 virus strains driving the epidemic within countries in Africa. The predominant mutation (90%) is the D614G substitution in the spike protein, and this aligns with those described in other settings around the globe. There were, however, foci of novel mutations, including deletions in the NSP12 protein, which is a target for antiviral drugs such as Remdesivir across the different countries from the various regions. Also, phylogenetic analyses show different clades that demonstrate the variability of the strains across countries in Africa, but these are substantially aligned to circulating strains from the countries of origin of the index cases of COVID-19 in the respective countries. The finding that the African SARS-CoV-2 spike proteins were highly identical to those of viruses from other geographical regions may provide a ray of hope on the cross efficacy of vaccines and therapeutics from other settings in Africa. However, mutations in critical proteins such as NSP12 may affect these interventions’ efficacy in African settings.

The entry of coronaviruses, including SARS-CoV and SARS-CoV-2, into host cells is essential for viral infectivity and pathogenicity^1113^. SARS-CoV-2 host cell entry is mediated by surface spike proteins recognizing and binding its human receptor angiotensin-converting enzyme – hACE2 via its Receptor binding domain (RBD) manner that is dependent on host cell pre-activation by host cell proprotein convertase furin^7^. Therefore, SARS-CoV-2 surface spike protein may affect the efficiency of viral spread. The majority of African spike proteins have a D614G substitution, as shown in the amino acid’s alignment. The D614G substitution was also identified to be predominant in Asian and European SARS-CoV-2 sequences^14^.

D614G substitution is a missense mutation (Aspartic acid (D) to Glycine (G) substitution), which predominantly emerged as a clade in Europe that eventually spread globally^15,16^. D614G substitution was one of eight missense mutations occurring in the S protein, and suggested to be relatively conserved, with potential effects on S protein interactions and functions^17^. The biological and clinical relevance of this mutation is linked to the type of amino acid and subsequent protein changes that arise. Studies using (*in silico)* structural bioinformatics to assess the effect of D614G mutation on SARS-CoV-2 virulence and epidemiology suggests that this mutation may be neutral to protein function^16^. Based on the structure of SARS-CoV-2 Spike protein (S) determined by Wrapp *et al*, using cryo-EM and 3-D modelling of the S-protein structure by Isabel *et al*, it was uncovered that this mutation leads to loss of four inter-chain destabilizing (hydrophobic-hydrophilic) contacts at a position relatively distal from the RBD with overall null effects on ACE2 and S protein interaction dynamics^16,17^. However, Yurkovetsky and colleagues showed that the D614G substitution, which is present in >97% of SARS-CoV-2 genomes globally, disrupts contact between S1 and S2 domains of the spike protein and significantly causes conformation shift^18^. Even though this mutation may have increased infectivity of SARS-CoV-2, it does not alter spike protein interactions, including with monoclonal antibodies targeting the S protein^18^.

The increase in the prevalence of D614G mutation across different geographical regions, as shown by Korber *et al*., maybe because this clade has a fitness advantage^19^. This variant of the SARS-CoV-2 virus was shown to grow to higher titers as pseudotyped virions, infect a broad range of human cell types such as lungs, liver and kidney and potentially have higher upper respiratory tract viral loads^19–21^.

The presence of multiple mutations, with the D614G mutation is the most predominant may be important for future studies. Specifically, the D614G substitution could potentially act as an anchor point for divergent S protein mutations especially within the African continent. Currently, efforts are geared toward the formulation of vaccines to combat the global scourge of the coronavirus pandemic. The target of the vaccines is the spike protein, which plays a central role in the viral entry into the host. The implications of the observed D614G mutation on the coverage of any vaccine that will be produced against the virus is not known. However, Ozono and Colleagues^22^, Hu and colleagues^23^ and Alexeev and Alekseyev^24^ several studies reported that antibodies generated due to natural infections of the two variants D614 or G614 showed cross neutralization re-activities thus indicating that the locus might not be within the epitope recognized by the host immune system and therefore not critical for humoral mediated immunity. As the D614G mutation does not prevent cross re-activities of antibodies generated against the variants, it is therefore very unlikely to interfere with the efficacy of any vaccine that will be eventually formulated. Therefore, we are of the courteous optimism that whichever vaccine that is finally approved will be beneficial to the African continent as the observed amino acid substitution is common to majority of the virus circulating in the different regions of the world apart from those from Wuhan, China.

The polymerization of SARS-CoV-2 depends on the main polymerase, the RNA-dependent RNA polymerase (RdRp, also named nsp12). Nsp12 is a 932-residue long enzyme which consists of two conserved domains, the nidovirus RdRp-associated nucleotidyltransferase (NiRAN) and the polymerase domains^25^. Using cryo-EM, Gao and colleagues determined the structure of SARS-CoV-2 Nsp12 bound to Nsp7 and Nsp8 which act as co-factors^26^. Upon comparison of sequences, SARS-CoV and SARS-CoV-2 sequences were found to be highly identical, including in their interactions with Nsps 7 and 8^25,27^. The finding of an amino acid deletion at position P314 (interface region of the enzyme) in both African and global consensuses compared to Wuhan consensus sequence may have significant effect on the interaction between nsp12 and Nsps 8 and 9. The implications of this deletion on viral fitness and its replicative capacity are not yet fully understood and therefore more work needs to be done especially to validate this deletion and how it affects viral fitness. Apart from the deletion in the NSP12, no substitution was observed. This has scientific validity as substantial substitutions in an enzymatic protein would have interfered with viral fitness and its replicative capacity.

Some Nsps have been the subject of clinical research into SARS-CoV-2 therapeutics. For instance, Remdesivir (GS-5734) is a nucleotide prodrug, a 1’ cyano-substituted adenosine analogue which has been shown to have broad antiviral activity including against coronaviruses MERS-CoV and SARS-CoV, by inhibiting CoV genome replication^28–30^. Specifically, NSP12 is a major target for the antiviral drug Remdesivir which plays a critical role in the replication of coronavirus genome. A mutation in the NSP12 of African SARS-CoV-2 genome may have critical ramifications for the ability of Africans to use drugs like Remdesivir and any NSP12 targeting drugs.

Genomes of viruses undergo evolution over time^24^. Our results showed that sequences originating from African countries clustered with each other, but overall African sequences did not form a monophyletic cluster. This lack of a monophyletic cluster may be because the analyzed sequences from the different African country were introduced from other countries. The fact that the genomes formed clades shows and suggests that at this time, there has been divergence of the SARS-CoV-2 from the continent to form new clades. This suggests that there may be multiple strains of SARS-CoV-2 virus circulating in the Africa continent. Further prospective studies are needed to monitor the evolution of the virus overtime both within the continent and globally.

As with all studies, this study has strengths and weaknesses. This study’s main strength is widespread the target African countries, cutting across all African continent (East, west, north, south and central). Secondly, this study focusses on a previously unstudied part of the world, as other studies have excluded analyzing the SARS-CoV-2 genomes in Africa. Finally, this study draws on available real-time data that continues to evolve and is a reasonable representation of the landscape at this time. On the other hand, the limitations include the low number of complete genomes sequenced and deposited as at that time. Research labs within the African continent seemed to be slow in gathering sequencing data in the initial months of the outbreak in Africa, which resulted in a relatively small number of complete genomes. Additionally, we cannot control for errors, lab-specific bias, handling bias, and sample collection bias because we are working with existing data collected by multiple labs. These limitations reduce the power of our results and inference. Finally, the findings in this study need to be validated using wet lab-based experiments.

## Conclusion

We conclude that the SARS-CoV-2 virus has shown genomic variability across countries in Africa, but most of the mutations that resulted in this variability align with those described in other countries of the world, including the index case in China. However, because these clades/strains are shown to predominantly cluster with strains from other continents, therapeutics (vaccines and drugs) designed based on SARS-CoV-2 strains in these continents could potentially be useful in Africa. However, the finding of novel mutations, though on a limited scale, may perhaps have implications on the manifestation, progression, severity and mortality of the infection with the virus. Our findings also call for region-specific testing of potential vaccines to ensure vaccines’ efficacy developed in other settings when used in Africa.

## Acknowledgement

MNL conceived and designed the project, while BCI carried out the bioinformatics analysis with MNL and OSE. The manuscript was written by MNL, BCI and OSE, with proofreading and additional inputs by SOD, EA, VTI, CIO, TO, AT, AM.

